# Suppression of meiotic crossovers in pericentromeric heterochromatin requires synaptonemal complex and meiotic recombination factors in *Drosophila melanogaster*

**DOI:** 10.1101/2024.12.19.629512

**Authors:** Nila M. Pazhayam, Sasha Sagar, Jeff Sekelsky

## Abstract

The centromere effect (CE) is a meiotic phenomenon that ensures meiotic crossover suppression in pericentromeric regions. Despite being a critical safeguard against nondisjunction, the mechanisms behind the CE remain unknown. Previous studies have shown that various regions of the *Drosophila* pericentromere, encompassing proximal euchromatin, beta and alpha heterochromatin, undergo varying levels of crossover suppression, raising the question of whether distinct mechanisms establish the CE in these different regions. To address this question, we asked whether different pericentromeric regions respond differently to mutations that impair various features that may play a role in the CE. In flies with a mutation that affects the synaptonemal complex (SC), a structure is hypothesized to have important roles in recombination and crossover patterning, we observed a significant redistribution of pericentromeric crossovers from proximal euchromatin towards beta heterochromatin but not alpha heterochromatin, indicating a role for the SC in suppressing crossovers in beta heterochromatin. In flies mutant for *mei-218* or *rec*, which encode components of a critical pro-crossover complex, there was a more extreme redistribution of pericentromeric crossovers towards both beta and alpha heterochromatin, suggesting an important role for these meiotic recombination factors in suppressing heterochromatic crossovers. Lastly, we mapped crossovers in flies mutant for *Su(var)3-9*. Although we expected a strong alleviation of crossover suppression in heterochromatic regions, no changes in pericentromeric crossover distribution were observed in this mutant, indicating that this vital heterochromatin factor is dispensable to prevent crossovers in heterochromatin. Our results indicate that the meiotic machinery plays a bigger role in suppressing crossovers than the chromatin state.

## Introduction

During the first meiotic division, recombination between homologous chromosomes is a crucial process that is required to promote their accurate segregation away from one another. Meiotic crossovers are a highly regulated phenomenon, with the meiotic cell tightly governing where along each chromosome crossovers can form. The rules that control crossover placement are commonly referred to as crossover patterning events and are an additional requirement in ensuring that homologs disjoin correctly during meiosis.

Of the various meiotic crossover patterning events that have been established (Sturtevant 1913; Beadle 1932; Owen 1950; Martini *et al*. 2006); reviewed in (Pazhayam *et al*. 2021)), the exclusion of crossovers near the centromere - commonly referred to as the centromere effect (CE) - occurs animals, fungi, and plants (Mahtani and Willard 1998; Copenhaver *et al*. 1999; Wu *et al*. 2003; Ghaffari *et al*. 2013; Vincenten *et al*. 2015; Nambiar and Smith 2016; Fernandes *et al*. 2024). Studies in both *Drosophila* and humans have shown a correlation between centromere-proximal crossovers and nondisjunction (Koehler *et al*. 1996; Lamb *et al*. 1996; Oliver *et al*. 2012).

Despite the importance of the CE in protecting against meiotic NDJ, little is known about how the CE is established or maintained. Studies that have looked at the *Drosophila* CE over the past century have largely attempted to establish the centromere or pericentromeric heterochromatin as being the final arbiter of crossover prevention in this region, but failed to reach a definitive conclusion (Mather 1939; Slatis 1955; Yamamoto and Miklos 1978; John 1985; Westphal and Reuter 2002; Mehrotra and Mckim 2006). Whether the CE is controlled by one primary mechanism of action, or several factors that must act together to suppress recombination in the region remains an unanswered question in the field, as does the identity and nature of these factors. Although the centromere effect has largely remained a mechanistic mystery since its discovery, certain modes of control have been ruled out in *D. melanogaster*. Disruption of centromere clustering, changes in centromere number, and changes in repetitive DNA dosage were shown to no *trans-*acting effects on the strength of the CE (PAZHAYAM *et al*. 2023).

The pericentromeric region in *Drosophila melanogaster*, as well as many other organisms including mammals, *Arabidopsis*, and fission yeast consists of a centromere embedded in large chunks of heterochromatinized repetitive DNA (Miklos and Cotsell 1990; Simon *et al*. 2015; Ghimire *et al*. 2024). Pericentromeric heterochromatin in *Drosophila* is heterogeneous (Figure 1), comprising two classes defined by sequence, staining patterns, and replication status. This is most clearly seen in polytene chromosomes, where the centromeres are embedded in regions that are densely staining and highly under-replicated, and the adjacent regions are more diffusely stained and are less under-replicated (Gall *et al*. 1971; Miklos and Cotsell 1990). The former, referred to as alpha heterochromatin, are composed largely of tandem arrays of highly repetitive satellite DNA sequences. The moderately stained regions, referred to as beta heterochromatin, are found between alpha heterochromatin and euchromatin, and have a high density of transposable elements interspersed within unique sequence. The unique sequences found in beta heterochromatin have made it possible to assemble much of it to the reference genome (Hoskins *et al*. 2015), whereas the alpha heterochromatin has not yet been assembled.

**Figure 1.**
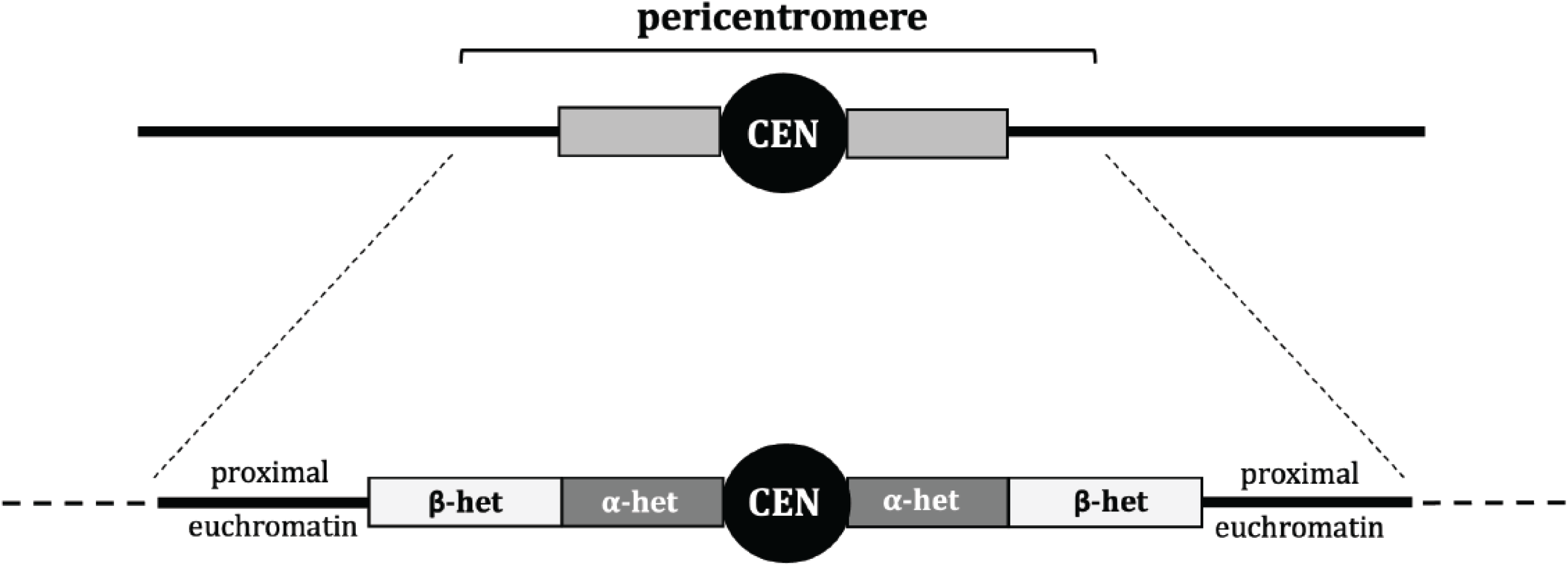
Schematic of the pericentromere region in *D. melanogaster*. Grey boxes indicate pericentromeric heterochromatin and thick black lines indicate euchromatin. In the lower image, the centromere indicated as CEN, alpha heterochromatin as α-het, and beta heterochromatin as β-het. Dashed lines indicate euchromatin that is not considered centromere-proximal and therefore excluded from our definition of the pericentromere.

These two classes of centromere-proximal heterochromatin also differ in crossover-suppression patterns. Hartmann et al. (2019) showed that in wild-type flies, meiotic crossovers are completely absent from alpha-heterochromatic regions, whereas crossover frequencies in beta heterochromatin and proximal euchromatin depend on distance from the centromere (Hartmann *et al*. 2019b). A similar pattern of centromere-proximal crossover suppression has been described in *Arabidopsis thaliana* (Fernandes *et al*. 2024), where the pericentromere is organized similarly to that *D. melanogaster*, with the centromere embedded in regions of highly repetitive heterochromatinized DNA that give way to less repetitive heterochromatinized DNA, followed by unique euchromatic sequence.

The existence of these two components of the CE raises the question of how they are established during meiosis, and whether distinct processes are responsible for their establishment and execution. It has been previously speculated that the “controlling systems” preventing crossovers in centromere-proximal euchromatin are different from those that prevent crossovers in pericentromeric heterochromatin (Carpenter and Baker 1982a; Szauter 1984), leading us to attempt to tease apart the mechanistic differences in proximal crossover suppression within the various regions of the pericentromere, including any - if they exist - between alpha and beta heterochromatin.

Evidence for centromere-proximal crossover suppression being a meiotically controlled phenomenon is abundant, and since the meiotic program is not a monolith, we focused on two facets: the synaptonemal complex (SC) and the proteins directing meiotic recombination. The SC is a protein structure that forms during meiosis between paired homologs and is the context within which meiotic recombination occurs. SC has been shown to be necessary for crossover formation as well as patterning in many species (Sym and Roeder 1994; Storlazzi *et al*. 1996; Page and Hawley 2001; Libuda *et al*. 2013; Voelkel-Meiman *et al*. 2015; Wang *et al*. 2015; Voelkel-Meiman *et al*. 2016; Billmyre *et al*. 2019). It has been proposed that the SC has liquid crystalline properties that helps mediate crossover designation and interference by providing a compartment within with the proteins that carry out these processes can diffuse (Morgan *et al*. 2021; Zhang *et al*. 2021; Von Diezmann *et al*. 2024). SC in pericentromeric heterochromatin has been reported to have morphological differences from the SC along euchromatin (Carpenter 1975). A 2019 study showed that the *Drosophila* SC component C(3)G plays a definitive role in suppressing pericentromeric crossovers (Billmyre *et al*. 2019). Collectively, these observations suggest the SC may have a crucial role in establishing the CE.

The second facet is the proteins that direct meiotic recombination. Hatkevich et al. (2017) showed that loss of Bloom syndrome helicase, an important DNA repair protein, lacked not only the CE, but also other forms of crossover patterning such as interference (Hatkevich *et al*. 2017). A 2018 study showed that the introduction of *D. mauritiana* orthologs of the pro-crossover genes *mei-217* and *mei-218* into *D. melanogaster mei-218* mutants attenuated crossover suppression around the centromere, as it is in *D. mauritiana*, suggesting that these genes mediate the strength of the CE in *D. melanogaster* (Brand et al. 2018). Mei-217 and Mei-218 are components of the meiotic-mini-chromosome-maintenance (mei-MCM) complex that is hypothesized the block the anti-crossover activity of Blm (Kohl *et al*. 2012) Analysis of the data of Hartmann *et al*. (2019) suggests that both *mei-218* and *rec*, which encodes the third component of the mei-MCM complex, may contribute to crossover suppression around the centromere. This, and data from other organisms showing genetic modes of suppressing pericentromeric crossovers through blocking or preventing Spo11-mediated meiotic DSBs (Vincenten *et al*. 2015; Nambiar and Smith 2018; Xue *et al*. 2018), suggests that the meiotic program is able to exert considerable control over the CE.

The heterochromatic nature of the pericentromere could also be a key factor contributing to the CE. Crossover suppression within heterochromatin as well as an effect of heterochromatin on crossover suppression in adjacent regions have previously been shown in *Drosophila* and other organisms (Slatis 1955; John 1985; Hartmann *et al*. 2019a; Fernandes *et al*. 2023; Fernandes *et al*. 2024). Westphal & Reuter (Westphal and Reuter 2002) observed elevated centromere-proximal crossovers in a several suppressor-of-variegation mutants that impact chromatin structure. Three of the *Su(var)* mutants in their study mapped to genes encoding proteins necessary for heterochromatin formation and maintenance, including HP1 (*Su(var)2-5*) and H3K9 methyltransferase (*Su(var)3-9*), as well as their accessory proteins (*Su(var)3-7*). Peng & Karpen (2009) showed that a hetero-allelic *Su(var)3-9* mutant had elevated DSBs in meiotic cells that colocalized with alpha-heterochromatic sequences, suggesting that *Su(var)3-9* is crucial to keeping DSBs out of alpha heterochromatin during meiosis. Together, these data suggest that the inherent heterochromatic nature of large portions of the pericentromere contributes to crossover suppression within it.

In this study, we measured centromere-proximal crossover frequencies, the strength of the CE, and crossover distribution patterns within different regions of the pericentromere: proximal euchromatin, beta heterochromatin, and alpha heterochromatin (Figure 1). We investigated three classes of mutants: structural (SC), meiotic, and heterochromatic. If multiple modes of crossover control are required to act in synchrony to suppress crossovers in centromere-proximal regions, we hypothesized that we would observe differences in where the CE is disrupted in each mutant class. The structural mutant we looked at was a *c(3)G* in-frame deletion mutant that leads to failure to maintain full-length SC by mid-pachytene (Billmyre *et al*. 2019). We observed significant CE defects on chromosome *2* in this mutant, along with a considerable redistribution of crossovers away from proximal euchromatin, towards beta but not alpha heterochromatin. This suggests that full length SC at mid-pachytene is required to suppress crossovers in beta heterochromatin. We also looked at mutants lacking *mei-218* and *rec*, which are crucial for crossover formation and patterning but have no known roles outside of meiosis/DNA repair (Carpenter and Baker 1982a; Hartmann *et al*. 2019a). Upon establishing that both mutants have a significantly weakened CE, we found a significant increase in heterochromatic crossovers in both beta and alpha heterochromatin at the expense of crossovers in proximal euchromatin. Surprisingly, the heterochromatic mutant in our study - *Su(var)3-9^null^ -* turned out to be dispensable not only for centromere-proximal crossover suppression, but also for preventing crossovers specifically in pericentromeric heterochromatin, as no significant redistribution of crossovers was observed between proximal euchromatin and pericentromeric heterochromatin. As *Su(var)3-9* is a gene crucial for heterochromatinization at the pericentromere (Schotta *et al*. 2002) and is also implicated in preventing meiotic crossovers in heterochromatin (Westphal and Reuter 2002), this result implies that chromatin-based steric hindrance/inaccessibility do not play as big of a role in keeping crossovers out of heterochromatic regions as various classes of meiotic factors necessary for crossover designation and patterning do.

Our results suggest that while the cell seems to require multiple facets of control to exclude crossovers in centromere-proximal regions during meiosis, the CE is a primarily meiotic phenomenon in *Drosophila*, with the meiotic program – both the structure providing the conduit for proteins that carry out recombination and the recombination proteins themselves – seemingly superseding heterochromatin in preventing heterochromatic crossovers.

## Results

### Synaptonemal complex protein C(3)G is necessary for centromere-proximal crossover suppression during meiosis

The synaptonemal complex is a protein structure that forms specifically between paired homologs during meiosis. In *Drosophila*, the SC is formed before meiotic DSBs are induced, and plays a crucial role in both DSB and crossover formation (Page and Hawley 2001; Mehrotra and Mckim 2006; Lake and Hawley 2012; Collins *et al*. 2014), as well as crossover patterning (Billmyre *et al*. 2019). To ask how important the *Drosophila* SC is in establishing the centromere effect, we measured recombination in a mutant defective for SC maintenance. *c(3)G^ccΔ2^* is a deletion the removes residues 346-361 from the coiled-coil domain of the transverse filament (Billmyre *et al*. 2019). This mutation results in loss of the SC structure by mid-pachytene. Interestingly, *c(3)G^ccΔ2^* flies display elevated centromere-proximal crossovers on chromosome *3*, which has a strong CE, but not on chromosome *X*, which has a weak CE, suggesting that C(3)G and a full-length SC are necessary to maintain a robust CE.

We asked whether C(3)G is important for pericentromeric crossover suppression on chromosome *2* as well by measuring crossover frequencies within a ∼40 Mb region that spans the centromere an includes euchromatin, beta heterochromatin, and alpha heterochromatin. Female flies heterozygous for markers on both arms of chromosome *2* were used to map recombination between the distal *2L* locus *net* and the proximal *2R* locus *cinnabar* (*cn*). The centromere on chromosome *2* lies in the interval between markers *purple* (*pr*) on *2L* and *cn* on *2R*, covering an approximate length of 20.5 Mb, including 11.2 Mb of assembled sequence and an estimated 4 Mb of alpha heterochromatin on *2L* and 5.3 Mb on *2R*.

Figure 2A shows crossover density along chromosome *2* (divided into five intervals by six recessive marker alleles) in wild-type flies and in *c(3)G^ccΔ2^* mutants. Total genetic length in this mutant is significantly increased in the mutant, from 48.05 cM in wild type to 64.01 cM (p<0.0001). While crossover distributions closely resemble wild-type in the three distal and medial intervals interval *2,* crossover frequencies in the interval spanning the centromere (*pr* - *cn*) and the adjacent interval (*b* - *pr*) are significantly increased in the *c(3)G^ccΔ2^* mutant (*p*<0.0001; Figure 2A). This suggests that chromosome *2,* like chromosome *3*, experiences a weaker centromere effect in this mutant.

**Figure 2.**
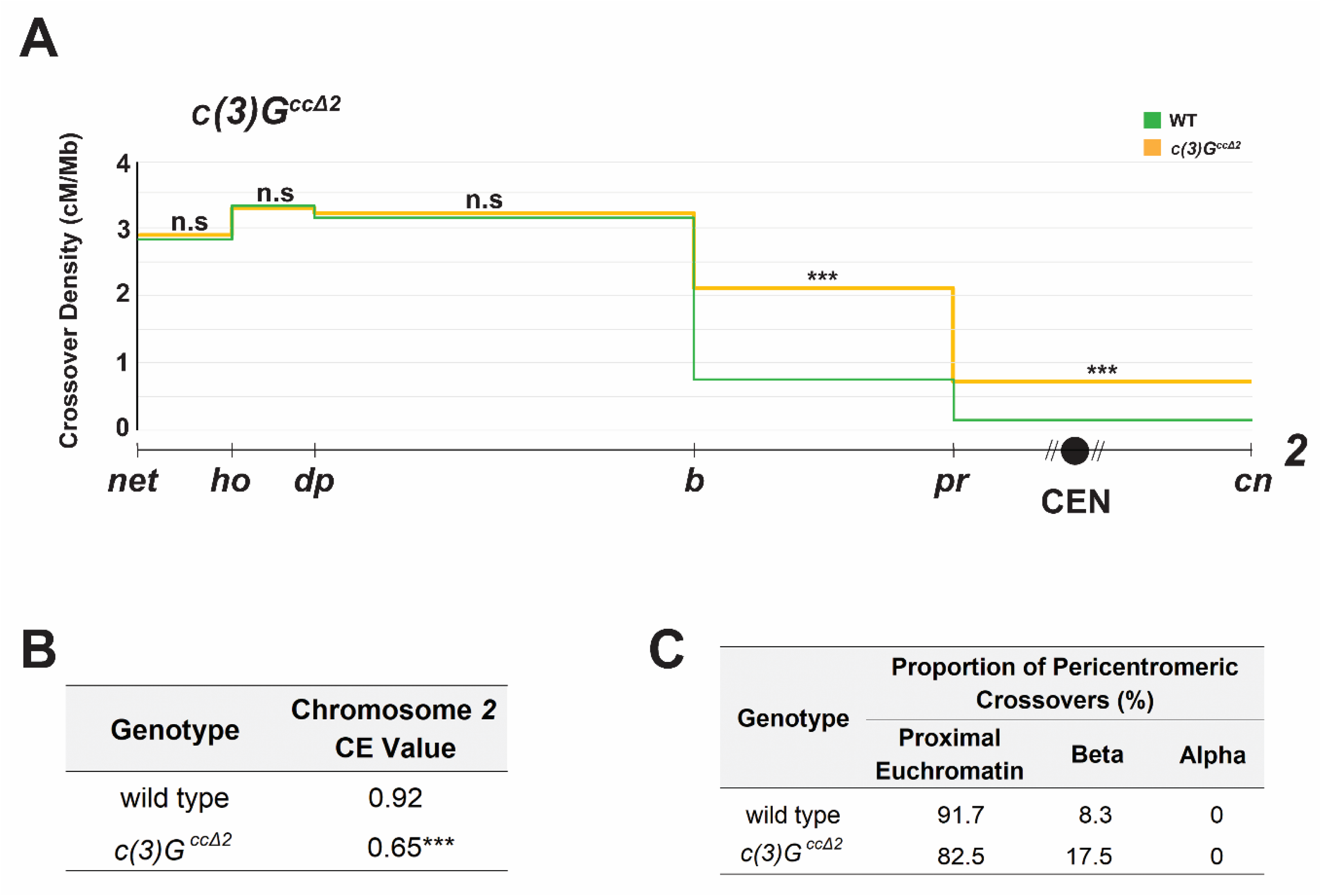
**A.** Crossovers in *c(3)G^ccΔ2^* (*n* = 5,918) and wild-type (*n* = 4,331) flies along chromosome *2* with the Y-axis indicating crossover density in cM/Mb and the X-axis indicating physical distances between recessive marker alleles that were used for recombination mapping. The chromosome *2* centromere is indicated by a black circle, unassembled pericentromeric repetitive DNA by diagonal lines next to it. A 2-tailed Fisher’s exact test was used to calculate statistical significance between mutant and wild-type numbers of total crossovers versus parentals in each interval. Complete dataset is in Supplementary Table S1. n.s *p* > 0.01, \**p* < 0.01, \*\**p* < 0.002, \*\*\**p* < 0.0002 after correction for multiple comparisons. **B.** Table showing CE values on chromosome *2* in wild type and *c(3)G^ccΔ2^* flies. \*\*\**p* < 0.0002. **C.** Table showing percentage of pericentromeric crossovers that occurred within each region of the pericentromere in wild type vs *c(3)G^ccΔ2^* mutant flies. Supplementary Figure S1 contains gel images of allele-specific PCRs for each SNP defining the boundaries of pericentromeric regions.

Since crossover frequencies measured in cM/Mb are based only on observed crossover numbers, we calculated a CE value that also takes into account crossover numbers expected if there no centromere-proximal suppression during meiotic recombination. This value considers crossover density in the centromeric interval as equal to the average density of the entire chromosome *2* region being studied and is a more biologically relevant measure of the CE as it is agnostic to differences in total crossover numbers between two genotypes.

WT flies have a CE value of 0.92 on chromosome *2* (PAZHAYAM et al. 2023), whereas the *c(3)G^ccΔ2^*mutant has a significantly lower CE value of 0.65 (*p*<0.0001; Figure 2B), consistent with a strong defect in the CE. This suggests that the maintenance of full-length SC throughout pachytene is essential for ensuring vigorous suppression of centromere-proximal meiotic crossovers in *Drosophila*.

### The synaptonemal complex protein C(3)G is necessary for crossover suppression in beta but not alpha heterochromatin

On observing that the *Drosophila* SC component C(3)G is crucial for centromere-proximal crossover suppression on chromosome *2,* we asked whether it plays a role in the distribution of crossovers across the various regions of the pericentromere. To determine this, we built flies of the desired mutant background that were heterozygous for isogenized *net-cn* and wild-type chromosomes. Through Illumina sequencing, we identified SNPs between these chromosomes, allowing us to fine map crossovers within the larger intervals defined by phenotypic markers. We collected every fly that had a crossover between *pr* and *cn* and, through allele-specific PCR, mapped the crossover to proximal euchromatin or beta heterochromatin on either arm, or to alpha heterochromatin. We defined beta heterochromatin as the region between where the H3K9me3 mark begins (Stutzman *et al*. 2024) and the most proximal SNPs on the current assembly (release 6.59 of the *D. melanogaster* reference genome). Alpha heterochromatin was defined as the region between the most proximal SNPs on *2L* and *2R*.

Intriguingly, the *c(3)G^ccΔ2^* mutant displayed a significant redistribution of crossovers across two of the three proximal regions. The distribution in this mutant, measured as percentages of total crossovers across the chromosomal region being studied, were significantly increased from wild type in proximal euchromatin and beta heterochromatin (Table 1). While only ∼2.7% of total crossovers on chromosome *2* form in proximal euchromatin in WT flies, *c(3)G^ccΔ2^*mutants had ∼4.1% of total chromosome *2* crossovers now found in this region (*p*=0.0012). Similarly, ∼0.9% crossovers in *c(3)G^ccΔ2^*mutants are found in beta heterochromatin, a significant (*p*=0.0002) increase from the ∼0.2% observed in wild-type flies (Table 1). Curiously, we observed no crossovers mapping to the region between our most proximal SNPs on *2L* and *2R,* meaning that no crossovers occurred in alpha heterochromatin, as in wild-type flies (Table 1). This suggests that while SC mutants are unable to maintain wild-type levels of crossover suppression in beta heterochromatin, they are as successful as wild-type flies in suppressing crossovers in alpha heterochromatin.

**Table 1.** Percentage of crossovers in the region of chromosome *2* being studied that occurred within each section of the pericentromere in wild type (WT) and mutants. ** *p*<0.01, *** *p*<0.001, **** *p*<0 0001. All others *p* >0.05.

| Genotype | Flies | Crossovers | Percentage of Chromosome 2 Crossovers |  |  |
| --- | --- | --- | --- | --- | --- |
|  |  |  | Proximal Euchromatin | Beta Heterochromatin | Alpha Heterochromatin |
| WT | 4331 | 2081 | 2.69 | 0.24 | 0 |
| *c(3)G<sup>ccΔ2</sup>* | 5918 | 3788 | 4.05** | 0.86*** | 0 |
| *mei-218<sup>null</sup>* | 12,339 | 284 | 10.21**** | 3.87**** | 0.35 |
| *rec<sup>null</sup>* | 16,776 | 848 | 10.97**** | 5.31**** | 0.94**** |
| *Su(var)3-9<sup>06/+</sup>* | 10,154 | 4871 | 2.24 | 0.16 | 0 |
| *Su(var)3-9<sup>null</sup>* | 8123 | 4289 | 2.98 | 0.37 | 0.02 |

We also calculated crossover frequencies in each region of the pericentromere as a percent of total pericentromeric crossovers in this mutant (Figure 2C), and observed a statistically significant redistribution from proximal euchromatin towards beta (*p*=0.0268) but not alpha heterochromatin (*p*=1.000), compared to WT.

Collectively, these data indicate that full length SC during mid-pachytene plays a role in maintaining wild-type levels of crossover suppression at the pericentromere (Figure 2A, 2B) as well as wild-type proportions of crossovers within proximal euchromatin and beta heterochromatin but is dispensable for crossover suppression within alpha heterochromatin (Figure 2C, Table 1).

### Meiotic recombination genes are necessary for centromere-proximal crossover suppression

Crossovers during meiosis are controlled by a meiotic program that designates and likely also patterns their formation along the length of the chromosome. To measure the influence of the meiotic program on centromere-proximal crossover suppression and the strength of the centromere effect, we first looked at a null mutant of the meiotic pro-crossover gene *mei-218*, which encode a component of the meiotic-mini-chromosome maintenance (mei-MCM) complex (Kohl *et al*. 2012). Mei-218 is crucial for the formation and patterning of meiotic crossovers (Baker and carpenter 1972; brand *et al*. 2018; Hartmann *et al*. 2019a). We addressed the role of *mei-218* in exerting the centromere effect by measuring recombination along chromosome *2*, between the markers *net* and *cinnabar*. Crossover density in *mei-218* null mutants is shown in Figure 3A. Consistent with its crucial role in crossover formation during meiosis, the *mei-218* mutant had a significantly reduced genetic length (2.30 cM, *p*<0.0001) along the chromosome *2* region being studied than wild-type flies did (48.05 cM). Notably, the distribution of crossovers along the chromosome in *mei-218* mutants appears to be almost flat, substantially different from the usual bell curve observed in wild-type flies. The genetic length of the interval containing the centromere was either close to or higher than crossover frequencies along the rest of the chromosome in this mutant, indicating an impaired centromere effect (Figure 3A).

**Figure 3.**
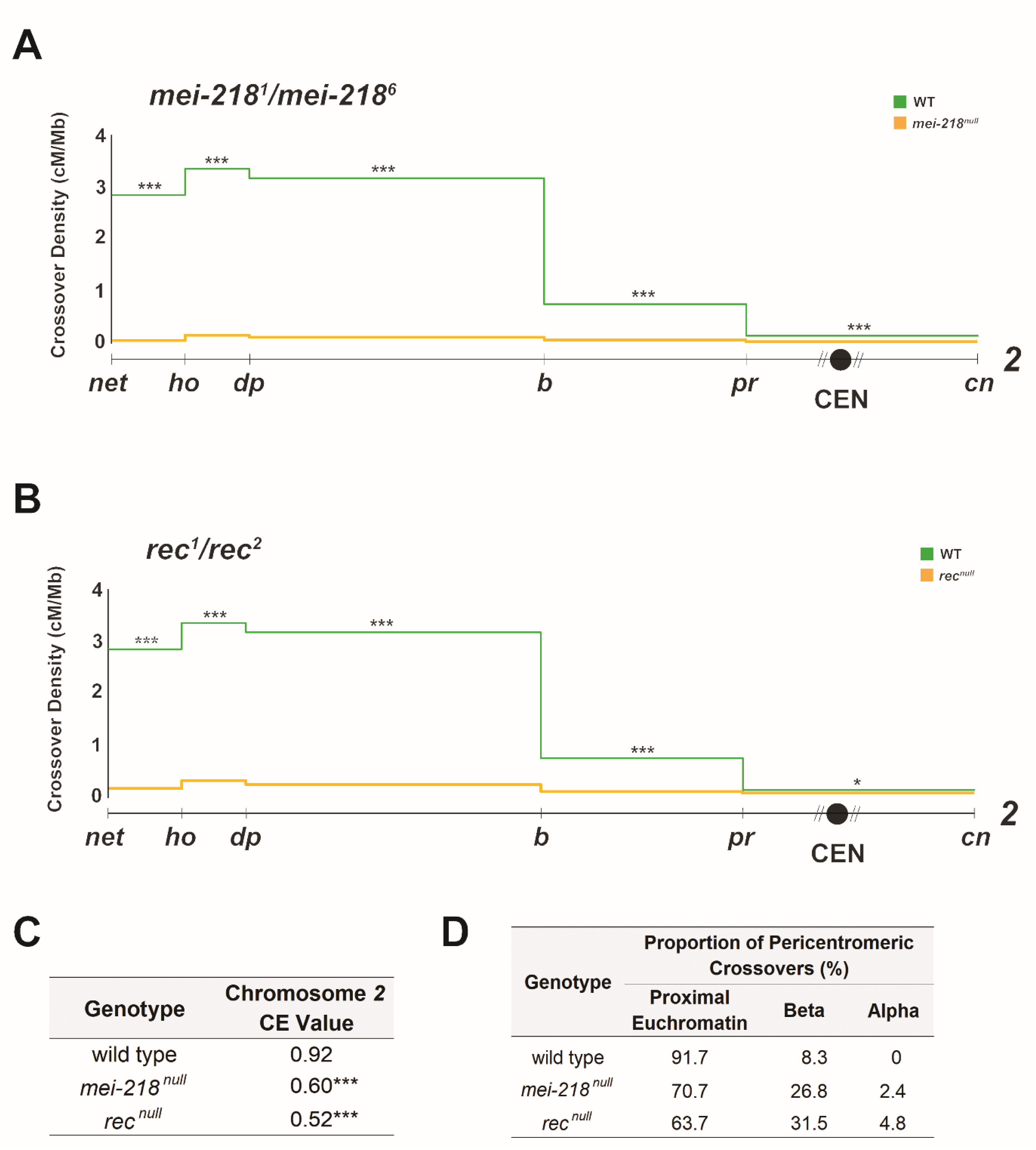
**A.** Crossovers in *mei-218^null^* (*n* = 12,339) and wild-type (*n* = 4,331) flies along chromosome *2* with the Y axis indicating crossover density in cM/Mb and the X axis indicating physical distances between recessive marker alleles that were used for recombination mapping. The chromosome *2* centromere is indicated by a black circle, unassembled pericentromeric repetitive DNA by diagonal lines. **B.** Crossovers in *rec^null^* (*n* = 16,776) and wild-type (*n* = 4,331). **C.** CE values on chromosome *2* in wild-type, *mei-218^null^*, and *rec^null^* flies. **D.** Table showing percentage of pericentromeric crossovers that occurred within each region of the pericentromere in WT, *mei-218^null^*, and *rec^null^* flies. For all panels, a 2-tailed Fisher’s exact test was used to calculate statistical significance between mutant and wild-type numbers of total recombinant versus non-recombinants in each interval (see Table S1 for complete datasets). n.s. *p* > 0.01, \**p* < 0.01, \*\**p* < 0.002, \*\*\**p* < 0.0002, after correction for multiple comparisons. Supplementary Figure S1 contains gel images of allele-specific PCRs for each SNP defining the boundaries of pericentromeric regions.

The *mei-218* mutant had a CE value of 0.60 on chromosome *2* (Figure 3C), a significant decrease from the WT chromosome *2* CE value of 0.92 (*p*<0.0001), further suggesting a very weak centromere effect in this mutant, consistent with what was observed by Hartman *et al*. (Hartmann *et al*. 2019a). Combined with the flat distribution of crossovers observed in this mutant, *mei-218* appears to be essential in establishing a robust suppression of crossovers near the centromere during meiosis.

To ask whether this importance in centromere-proximal crossover suppression extended to other pro-crossover meiotic genes, we also studied mutants defective for *rec*, which encodes another mei-MCM component (Kohl *et al*. 2012). Figure 3B shows crossover density along chromosome *2* in *rec* null mutants, which also show a significant decrease in genetic length (5.05 cM; *p*<0.0001) from the wild-type level. Crossovers in this mutant followed the pattern of the *mei-218* mutant, with a much flatter distribution observed along the chromosome than in wild-type flies. The genetic length of the interval spanning the centromere was once again higher than or much closer to the genetic lengths of intervals in the middle of the chromosome arm, suggesting that *rec* mutants also have a diminished centromere effect. This is further corroborated by the CE value of *rec* mutant flies (0.52), significantly reduced from WT chromosome *2 C*E value of 0.92 (p<0.0001) (Figure 3C), indicating that Rec is also crucial for maintaining a strong centromere effect. Overall, these results demonstrate that genes encoding two components of the mei-MCM complex - *mei-218* and *rec* - are independently necessary to ensure that crossovers form at the right frequencies, and to guarantee centromere-proximal crossover suppression in *Drosophila*.

### Meiotic recombination genes are necessary for crossover suppression in alpha and beta heterochromatin

On observing that the meiotic mutants *rec* and *mei-218* both have an ablated CE, we asked whether these genes are also necessary to maintain wild-type patterns of crossover distribution within the pericentromere. Hartmann *et al*. (2019b) previously fine mapped centromere-proximal crossovers in *Blm* mutants, which also lack a functional CE, and observed a flat crossover distribution that extended into proximal euchromatin and beta heterochromatin, but never into alpha heterochromatin. They concluded that Blm is necessary to maintain the distance-dependent CE observed in beta heterochromatin and proximal euchromatin, but that the complete suppression of crossovers observed in alpha heterochromatin is likely due to the region not being under genetic/meiotic control, hypothesizing instead that highly repetitive regions do not experiencing meiotic DSBs.

This pattern of crossover redistribution in *Blm* mutants is similar to what we observed in the SC mutant *c(3)G^ccΔ2^* is consistent with an important contribution of the SC in regulating meiotic recombination. Since the CE in both *rec* and *mei-218* mutants is weakened much like in *Blm* and *c(3)G^ccΔ2^* mutants, we sought to ask if fine mapping crossovers within the pericentromere in *mei-218* and *rec* mutants would reveal the same patterns of crossover redistribution observed in *Blm^null^* and *c(3)G^ccΔ2^*flies. Surprisingly, pericentromeric crossover distribution patterns in the *mei-218* and *rec* mutants were different from both *Blm* and *c(3)G^ccΔ2^*mutants. In *mei-218* mutants, 10.2% of total chromosome *2* crossovers were within proximal euchromatin, a significant increase from both the WT value of 2.7% in this region, as well as the *c(3)G^ccΔ2^* value of 4.05% (*p*<0.0001 for both comparisons). Similarly, 3.9% of total crossovers in *mei-218* mutants form in beta heterochromatin, also a significant increase compared to wild-type (p<0.0001) and *c(3)G^ccΔ2^* (p=0.0002) flies (Table 1).

Interestingly, we observed an increase in crossover frequencies in the region described as alpha heterochromatin, with 0.4% of total chromosome *2* crossovers in *mei-218* mutants forming between our most proximal SNPs, compared to none in both wild-type and SC mutant flies (Table 1). The increase isn’t statistically significant (*p* = 0.35), but statistical power is limited by the severe reduction in total crossovers in *mei-218* leading to few pericentromeric crossovers (41 from >12,000 flies scored). Because we never saw a crossover between the most proximal SNPs in wild type (n=132), the increase observed in the *mei-218* mutant may be biologically relevant.

We then looked at pericentromeric crossover distributions in the *rec* mutant and observed similar patterns to those of the *mei-218* mutant. When compared to wild type, crossover frequencies, measured as a percent of total crossovers across chromosome *2*, were increased in all three regions of *rec* mutants (Table 1). crossover frequencies increased to ∼11% in proximal euchromatin, ∼5.3% in beta heterochromatin, and ∼0.9% in alpha heterochromatin, all significant (*p*<0.0001) changes from crossover frequencies in the respective pericentromeric regions of wild-type and SC mutant flies.

We also calculated crossover frequencies as a percent of total pericentromeric crossovers (Figure 3D) and observed a statistically significant redistribution from proximal euchromatin towards beta heterochromatin in both *mei-218* (*p*=0.0049) and *rec* (*p*<0.0001) mutants, compared to wild-type flies. Compared to *c(3)G^ccΔ2^*flies, *mei-218* mutants did not exhibit a significant redistribution of crossovers from proximal euchromatin to beta heterochromatin (*p*=0.1824), but *rec* mutants did (*p*=0.0032). *rec* mutant flies also displayed a highly significant redistribution of pericentromeric crossovers from proximal euchromatic regions towards alpha heterochromatin, compared to both wild-type (*p*=0.0016) and SC mutant (*p*=0.0008) flies.

Collectively, these results suggest that when the mei-MCM complex is lost, there is a significant repositioning of crossovers within the pericentromere, compared to both wild type and the SC mutant in our study. More specifically, we observe a clear redistribution of pericentromeric crossovers away from proximal euchromatin and into both alpha and beta heterochromatin. Centromere-proximal crossovers in both mutants can reach further into pericentromeric heterochromatin than in wild-type, *Blm* mutant, or SC mutant flies, indicating not only a weakening of the strength of the CE but also its reach along the chromosome. This is particularly striking, as heterochromatic crossover suppression has been widely thought to happen through non-meiotic mechanisms (Carpenter and Baker 1982b; Szauter 1984; Westphal and Reuter 2002; Mehrotra and Mckim 2006), possibly through heterochromatinization and steric hindrances to DSB and recombination machinery. We had expected to see increases in crossovers within pericentromeric heterochromatin only in mutants of important heterochromatin genes. Instead, crossovers within heterochromatin seem to unambiguously be under meiotic control.

### *Su(var)3-9* is dispensable for centromere-proximal crossover suppression during meiosis

On observing that the meiotic machinery – in the form of both SC and recombination proteins – is necessary to prevent heterochromatic crossovers, we asked what pericentromeric crossover distributions look like in a heterochromatin mutant. As the majority of the chromosomal region described as the pericentromere is heterochromatic, we wanted to investigate whether mutations in genes necessary for heterochromatin formation and maintenance disrupt the CE and/or the suppression of heterochromatic crossovers to even greater extents than observed in our SC and meiotic recombination mutants.

To this end, we wished to look at a some of the suppressor of variegation mutants that were reported to have elevated centromere-proximal crossovers (Westphal and Reuter 2002). Of the genes in that study, *Su(var)3-7* and *Su(var)3-9* were of the most interest to us, as they encode critical heterochromatin-associated proteins. *Su(var)3-9* codes for the H3K9 methyltransferase responsible for methylating pericentromeric heterochromatin, and SU(VAR)3-7 functions as an HP1 companion (ClÉard *et al*. 1997; Delattre *et al*. 2000) and potential anchor for the HP1 and SU(VAR)3-9 complex (Westphal and Reuter 2002).

We hypothesized that the elevation of pericentromeric crossovers observed on chromosome *3* in the *Su(var)3-7* heterozygote and the *Su(var)3-7 Su(var)3-9* double heterozygote in (Westphal and Reuter 2002) would hold true on chromosome *2,* and that the excess centromere-proximal crossovers in these mutants would map to the heterochromatic regions of the pericentromere. We assayed flies with a heteroallelic *Su(var)3-9* genotype previously observed to have elevated DSBs in female meiotic cells (Peng and Karpen 2009). We hypothesized that this elevation would lead to an increase in centromere-proximal crossovers and a subsequent weakening of the centromere effect.

When crossover distribution was measured along chromosome *2* in *Su(var)3-9^06^/ Su(var)3-9^17^* females, we found in increase in genetic length in the region being studied, from 48.05 cM in wild-type females to 52.8 cM in the mutant (*p*=0.0041); however, this elevation in genetic length comes from an increase in distal, euchromatic crossovers that lie outside of the purview of SU(VAR)3-9’s H3K9 methylation functions. Furthermore, crossover frequencies within the interval containing the centromere were not different from wild-type levels, and no change in crossover density was observed (Figure 4A). The chromosome *2* CE value in this mutant (0.91) was also unchanged from the WT chromosome *2* CE value (0.92) (Figure 4C), further indicating that the centromere effect remains intact. This is despite the reported elevation in DSBs in meiotic cells in this mutant (Peng and Karpen 2009). This suggests that crossover homeostasis is intact in this mutant, consistent with meiotic cells employing multiple levels of control to ensure crossover suppression around the centromere.

**Figure 4.**
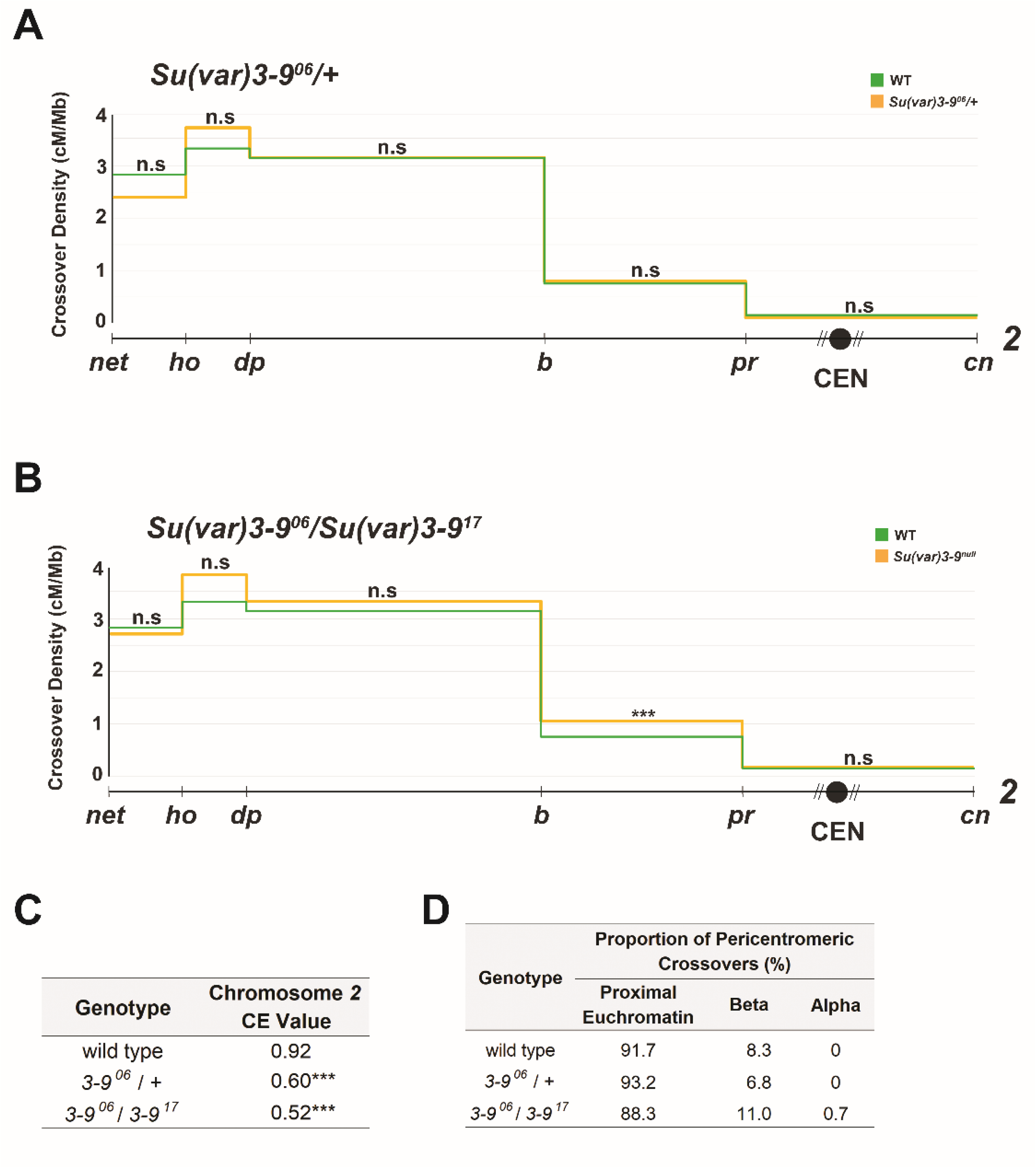
**A.** Crossovers in *Su(var)3-9^06^*/+ (*n* = 10,154) and wild-type (*n* = 4,331) flies along chromosome *2* with the Y axis indicating crossover density in cM/Mb and the X axis indicating physical distances between recessive marker alleles that were used for recombination mapping. The chromosome *2* centromere is indicated by a black circle, unassembled pericentromeric DNA by diagonal lines. **B.** Crossovers in *Su(var)3-9^06^/Su(var)3-9^17^* (*n* = 8,123) and wild-type (*n* = 4,331) flies. **C.** CE values on chromosome *2* in WT, *Su(var)3-9^06^*/+, and *Su(var)3-9^06^/Su(var)3-9^17^* flies. **D.** Percentages of pericentromeric crossovers that occurred within each region of the pericentromere in wild-type, *Su(var)3-9^06^*/+, and *Su(var)3-9^06^/Su(var)3-9^17^* flies. For all panels, a 2-tailed Fisher’s exact test was used to calculate statistical significance between mutant and wild-type numbers of total crossovers versus non-recombinants in each interval. n.s *p* > 0.01, \**p* < 0.01, \*\**p* < 0.002, \*\*\**p* < 0.0002 after correction for multiple comparisons. Supplementary Table S1 contains complete datasets. Supplementary Figure S1 contains gel images of allele-specific PCRs for each SNP defining the boundaries of pericentromeric regions.

We also measured crossover distribution along chromosome *2* in a *Su(var)3-9^06^/+* heterozygote (Figure 4B) and observed no changes from wild type in total genetic length (47.97 cM) or in crossover density in the centromeric interval. The *Su(var)3-9^06^* heterozygote had a CE value of 0.93 (Figure 4C), not significantly different from the wild-type CE value of 0.92 (*p*=0.2050), indicating that the centromere effect remains robust in this mutant.

Collectively, these results demonstrate that the H3K9 methyltransferase necessary for heterochromatinization of pericentromeres is dispensable both for the formation of crossovers and for suppression of crossovers in pericentromeric regions. Crossover homeostasis and CE machinery are reliably able to function in these mutants to guarantee that crossovers form at the correct frequencies and in the right chromosomal regions.

### *Su(var)3-9* is dispensable for suppressing crossovers in heterochromatin

Although no changes were observed in the strength of the CE in *Su(var)3-9* mutants, it is still possible that crossover distribution within the pericentromeric interval is affected. Peng & Karpen (2009) reported in 2009 that many of the excess DSBs they observed in meiotic cells of *Su(var)3-9^06^/Su(var)3-9^17^* mutants co-localized with signals from fluorescent *in situ* hybridization of probes to satellite DNA sequences, something never seen in wild-type flies. This suggests that there may be a redistribution of crossovers within the pericentromeric interval towards alpha-heterochromatic regions. However, when we measured crossover frequencies in the *Su(var)3-9^06^/Su(var)3-9^17^* mutant in each of the pericentromeric regions (as a percent of total crossovers across the chromosomal region being studied) we found that they closely resembled WT levels (Table 1), with ∼3% of total crossovers on chromosome *2* forming in proximal euchromatin and ∼0.4% forming in beta heterochromatin. These are not significant changes from wild-type percentages (*p*=0.4406 and 0.3363, respectively).

We also calculated crossover frequencies within each pericentromeric region as a percent of total crossovers within the pericentromere, and once again observed no significant changes from wild-type frequencies, with 88% of pericentromeric crossovers mapping to proximal euchromatin (*p*=0.8614 compared to wild type) and 11% to beta heterochromatin (*p* = 0.5486) (Figure 4D). However, we did observe one crossover between the most proximal SNPs, which we never saw in our dataset from wild-type females.

We also looked at pericentromeric crossover distributions in the *Su(var)3-9* heterozygote tested by Westphal and Reuter (2002) (Westphal and Reuter 2002), but saw no significant changes in total or pericentromeric crossover frequencies in proximal euchromatin, beta heterochromatin, or alpha heterochromatin. Similar to wild-type flies, 2.2% of total crossovers in this mutant were in proximal euchromatin, 0.2% were in beta heterochromatin, and 0% were in alpha heterochromatin (Table 1). Percentages of total pericentromeric crossovers also closely resembled wild-type percentages, with 93.2% occurring in proximal euchromatin and 6.8% occurring in beta heterochromatin (Figure 4D).

Overall, the lack of any significant redistribution of crossovers within the pericentromere tells us that meiosis successfully able to suppress pericentromeric crossovers in *Su(var)3-9^06^/Su(var)3-9^1^7* mutants. Peng & Karpen (2007) showed that this mutant has reduced H3K9 methylation at repetitive regions of the genome, suggesting that H3K9 methylation – a hallmark of heterochromatinization – within the pericentromere is surprisingly dispensable for crossover suppression in beta heterochromatin and for keeping pericentric crossovers within proximal euchromatin. It also appears to be largely or completely dispensable for crossover suppression in alpha heterochromatin. Despite allowing for more heterochromatic DSBs during meiosis, the *Su(var)3-9* mutant can maintain wild-type distributions of crossovers within the *Drosophila* pericentromere, completely unlike the SC and meiotic recombination mutants in our study.

## Discussion

Previous studies have shown that the centromere effect manifests differently in different regions of the pericentromere, with alpha heterochromatin displaying no crossovers and beta heterochromatin and proximal euchromatin displaying crossover suppression that diminishes with increasing distance from the centromere (Hartmann *et al*. 2019b; Fernandes *et al*. 2024). This suggests that the CE may be established via distinct mechanisms in different pericentromeric regions, motivating us to look at patterns of centromere-proximal crossover formation in three classes of mutants. These mutants affect either SC maintenance (Billmyre *et al*. 2019), meiotic recombination (Baker and carpenter 1972; hartmann *et al*. 2019a), or heterochromatin formation (Schotta *et al*. 2002), and were utilized to ask whether each of these processes exerts control over crossover suppression in independent regions of the pericentromere.

Our data show that crossover regulation at the pericentromere is indeed multi-faceted, with each class of mutants exhibiting distinct patterns of crossover formation in the various pericentromeric regions, summarized in Figure 5. We discuss the mechanistic implications of these results below.

**Figure 5.**
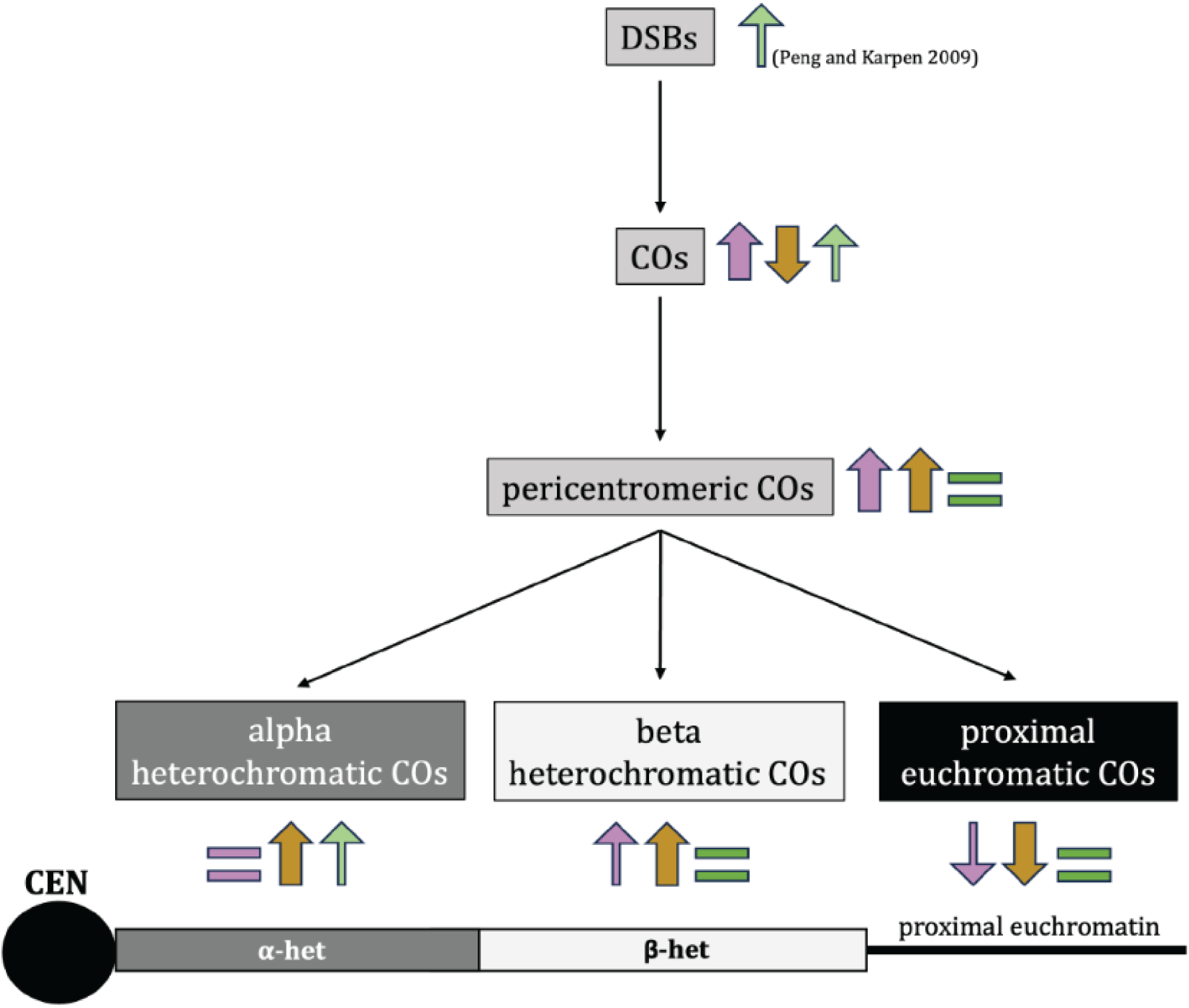
Summary of the effects of each mutant in this study on the formation of DSBs, crossovers, pericentromeric crossovers, alpha-heterochromatic crossovers, beta-heterochromatic crossovers, and proximal euchromatic crossovers. The arrows indicate whether there is an increase or decrease in the indicated event, with colors denoting the mutant in question. Purple is *c(3)G^ccΔ2^,* dark yellow is *mei-218^null^* and *rec^null^* combined, green is *Su(var)3-9^06^/Su(var)3-9^17^.* Thickness of the arrows and intensity of color indicate strength of the increase/decrease. A schematic of a telocentric chromosome is shown below, with the centromere, alpha heterochromatin, beta-heterochromatin, and proximal euchromatin indicated.

### Synaptonemal complex and the centromere effect

The SC is a meiotic structure essential for recombination in *Drosophila*, likely through facilitating the movement of meiotic recombination factors - such as the mei-MCM complex – along chromosomes. It provides a framework of sorts for the process of crossing-over and has been shown to contribute towards crossover patterning in various ways (Sym and Roeder 1994; Wang *et al*. 2015; Billmyre *et al*. 2019; Zhang *et al*. 2021). We sought to ask how disrupting it would affect pericentromeric crossover suppression and distribution.

The SC mutant in our study is an in-frame deletion of the SC gene *c(3)*, which encodes the transverse filament of the *Drosophila* SC and is essential for SC assembly as well as meiotic recombination (Page and Hawley 2001). The allele we used – *c(3)G^ccΔ2^* – has defects in SC maintenance and fails to retain its full length structure by mid-pachytene (Billmyre *et al*. 2019). This mutant was also shown to exhibit increased centromere-proximal crossovers on chromosome *3,* making it an ideal candidate to test how the SC contributes to the CE as well as to suppressing crossovers in different regions of the pericentromere.

Our data show the *c(3)G* mutant having a significantly weaker CE (Figure 2A, 2B) as well as a pericentromeric crossover redistribution phenotype that is intermediate between our meiotic recombination mutants and wild-type flies. While a significant increase in percentage of total crossovers is observed in both proximal euchromatin and beta heterochromatin in *c(3)G^ccΔ2^*flies, no change is observed in alpha-heterochromatic crossover frequencies when compared to wild type (Table 1). Additionally, the increases observed in proximal euchromatin and beta heterochromatin in the SC mutant do not reach the levels observed in either meiotic mutant (Table 1, Figure 2C, Figure 3C), indicating that while full length SC during mid-pachytene is necessary for centromere-proximal crossover suppression and to maintain wild-type proportions of crossovers within proximal euchromatin and beta heterochromatin, it doesn’t appear to be as crucial as the meiotic-MCM genes.

This is surprising as it tells us that despite *c(3)G^ccΔ2^* mutants having an ablated CE, meiotic cells in this mutant are still able to regulate crossover formation within the pericentromere and prevent the spread of excess centromere-proximal crossovers into alpha heterochromatin, and even into beta heterochromatin at the levels allowed in *mei-218* and *rec* mutants. Like Blm, C(3)G appears to be necessary to maintain the distance-dependent CE observed in beta heterochromatin and proximal euchromatin, but dispensable for the complete suppression observed in alpha heterochromatin. These data suggest that it is possible to disrupt the CE in different ways – using different classes of mutants – that may allow an increase in crossovers within one region of the pericentromere but not another, or even different levels of crossover increases within the same region.

Our observations also fit well with the SC serving as a conduit for the recombination proteins that designate and pattern crossovers during prophase I (Rog *et al*. 2017; Zhang *et al*. 2021; Fozard *et al*. 2023; Von Diezmann *et al*. 2024). Without any SC, as in the case of *c(3)G* null mutants, flies are completely unable to make meiotic crossovers (Page and Hawley 2001). This could be because meiotic proteins now lack a phase through which to travel along the length of paired homologs. In the *c(3)G^ccΔ2^* mutant, however, crossovers still form – at rates even higher than in wild type – but the CE is drastically weakened, which suggests that meiotic proteins can diffuse enough to designate crossovers along the chromosome, but somehow lose the ability to suppress them at the pericentromere. One explanation for this could be that centromere-proximal crossover suppression might be enforced after initial crossover designation. The *c(3)G^ccΔ2^* mutant has full length SC in early and early/mid-pachytene, but this is lost by mid-pachytene. It is possible that initial crossover designation occurs in early-pachytene, but the CE is established in mid-pachytene, and therefore severely disrupted in this mutant. Crossover distribution patterns being altered in *c(3)G^ccΔ2^* flies could also be related to timing, as it is possible that crossover suppression in alpha heterochromatin happens early, when the SC in these mutants is still fully intact, with beta-heterochromatic and proximal euchromatic crossovers being suppressed at mid-pachytene or later, when full length SC is lost in the mutant. Measuring the strength of the CE as well as pericentromeric crossover patterns in the other deletion mutants described in (Billmyre *et al*. 2019) that lose full length SC at different times during pachytene could shed light on which ones are important for crossover suppression in the different pericentromeric regions.

An interesting point to note about the *c(3)G^ccΔ2^* mutant is that while it has a weaker than wild-type CE on chromosomes *2* and *3*, the weak CE on the *X* chromosome appears not to be affected (Billmyre *et al*. 2019). Curiously, another *c(3)G* deletion described by Billmyre *et al*. (2019) – *c(3)G^ccΔ1^*– displays CE defects on all three chromosomes, suggesting that different aspects of SC function and maintenance are important for CE establishment on different chromosomes. This suggests that CE mechanism may not be uniform across the genome. Investigating how pericentromeric crossover distributions are changed in *c(3)G* mutants that have an ablated CE on all three chromosomes may illuminate which aspects of SC function are important across the board, and which are important only for certain chromosomes.

### Recombination machinery and the CE

The recombination genes in our study – *mei-218* and *rec -* encode two major components of the mei-MCM complex, a pro-crossover protein complex necessary for both crossover formation and patterning during meiosis (Kohl et al. 2012). As these proteins are crucial for meiotic recombination but have no SC defects (Carpenter 1979), they provide data that is easily separable from the *c(3)G^ccΔ2^* mutant, allowing us to draw conclusions about the importance of recombination machinery independently of SC-mediated influences to centromere-proximal crossover suppression.

Based on data from the SC mutant in our study, as well as *Blm* mutants (Hatkevich *et al*. 2017), we hypothesized that *mei-218* and *rec* mutants would exhibit a similarly defective CE, with increased pericentromeric crossovers in proximal euchromatin and beta heterochromatin but no changes from the complete crossover suppression in alpha heterochromatin. While we did observe significantly weaker centromere effects in both recombination mutants, we were surprised to see a substantial increase of total crossover percentages across all three regions of the pericentromere, with a significant redistribution of crossovers away from proximal euchromatin towards both beta and alpha heterochromatin (Table 1; Figure 3D). It must be noted here that the current assembly of the *Drosophila* reference genome is incomplete, and that the crossovers we recover in what we call alpha heterochromatin – defined as the region between the most proximal SNPs in our study – may still be occurring within beta heterochromatin. Nevertheless, *mei-218* and *rec* mutants having any crossovers between our most proximal SNPs is noteworthy, as none were ever observed in *Blm* or *c(3)G* mutants (Hartmann *et al*. 2019; Figure 3D). This suggests that the mei-MCM complex suppresses crossovers deeper into beta heterochromatin and/or alpha heterochromatin than Blm or SC. These data also indicate that these two parts of the meiotic recombination machinery may have distinct areas of control within the pericentromere. Pericentromeric crossover distributions in double mutants could shed light on whether Blm and the mei-MCM complex work in tandem to maintain the CE and are equally important to suppress crossovers in the region.

Aside from how crossover distribution in these mutants differs from the *Blm* and *c(3)G* mutant, it is also unexpected and noteworthy that Mei-218 and Rec are necessary to prevent crossovers in heterochromatin. Previous data has shown that while “recombination-defective meiotic mutants” such as *mei-218* can change euchromatic crossover distribution patterns on chromosome *X* and, unexpectedly, *4*, they do not allow for the formation of heterochromatic crossovers on either chromosome (Sandler and Szauter 1978; Carpenter and Baker 1982b). Szauter (1984) inferred that the mechanisms “that prevent crossovers in heterochromatin are distinct from those that specify the distribution of crossovers in the euchromatin” (Szauter 1984). Our chromosome *2* results appears to contradict these conclusions, showing not only that heterochromatic crossovers *can* be under the control of meiotic machinery in *Drosophila*, but also reinforcing our hypothesis that the CE is mediated differently on different chromosomes.

### Heterochromatin and the centromere effect

While both facets of the meiotic machinery tested in our study – SC and recombination genes – were observed to suppress heterochromatic crossovers, we wondered whether a stronger influence on pericentromeric crossover suppression is exerted by genes essential for heterochromatin formation, given that much of the pericentromere is heterochromatic. To test this, we used mutants of *Su(var)3-9*, the H3K9 methyltransferase that methylates and aids in the heterochromatinization of the pericentromere. Specifically, we tested a *Su(var)3-9* heterozygote - *Su(var)3-9^06^/+ -* as well as a heteroallelic null mutant *Su(var)3-9^06^/Su(var)3-9^17^*that was previously shown to have elevated DSBs within alpha heterochromatin in meiotic cells (Peng and Karpen 2009). Hypothesizing that heterochromatic crossover suppression is primarily chromatin-based, we expected to see a significantly greater number of crossovers in both heterochromatic regions of the pericentromere in this mutant compared to wild-type and to both classes of meiotic mutants. Surprisingly, we saw no change from wild type in CE value or total crossover distribution patterns in proximal euchromatin or beta heterochromatin, suggesting that pericentromeric crossover suppression is not mediated by this H3K9 methyltransferase, despite it being a key component of pericentromeric heterochromatinization. It appears that heterochromatic crossovers are not suppressed during meiosis because they occur in heterochromatin and may be subject to steric hindrances, but by virtue of them being under control of meiotic machinery.

Interestingly, we did recover one crossover between our most proximal SNPs in the *Su(var)3-9^06^/Su(var)3-9^17^* mutant. We believe this could be biologically relevant, as we observe complete suppression of crossovers in this region in wild-type flies. While this one crossover may be in unassembled beta heterochromatin, it is notable that *Su(var)3-9^06^/Su(var)3-9^17^* do not exhibit increased crossovers in beta heterochromatin. It is possible that this crossover was mitotic in origin. Among 3393 progeny of *Su(var)3-9^06^/Su(var)3-9^17^*males, which do not have meiotic recombination, we recovered a single crossover, in beta heterochromatin (Supplemental Table S1). Mitotic crossovers in the male germline are extremely rare in wild-type males (McVey *et al*. 2007), so this may indicate a true increase in these mutants. We note that the elevated DSBs in female meiotic cells reported by Peng and Karpen (2009) may not behave like typical meiotic DSBs in terms of repair mechanisms and regulation.

### Conclusions

Our study demonstrates that crossover control at the *Drosophila* pericentromere is multifaceted, and that a collaborative effort between diverse factors that include the SC, various recombination proteins, and even chromatin state may be necessary to establish or enforce the centromere effect. We show that suppression of meiotic crossovers within heterochromatin appears to be influenced less, if at all, by the chromatin state and more by the meiotic machinery. Our data, in conjunction with studies from other labs, suggests that the mechanisms behind the centromere effect may vary among chromosomes, providing fertile ground for future research on pericentromeric crossover suppression in *Drosophila* and other species.

## Materials and Methods

### Fly stocks

Flies were maintained at 25 C on a corn meal-agar medium. The *Oregon-R* stock used as our wild-type control was generously provided by Dr. Scott Hawley. The *mei-218* mutant alleles used in this study (*mei-218^1^* and *mei-218^6^*) are described in (Baker and carpenter 1972; mckim *et al*. 1996). The *rec* mutant alleles used in this study (*rec^1^* and *rec^2^*) are described in (Grell 1978; Matsubayashi and Yamamoto 2003; Blanton *et al*. 2005). The *y ; Su(var)3-9^06^/TM3 Sr* and *y ; Su(var)3-9^17^/TM3 Sr* stocks were generously provided by Dr. Gary Karpen. The *y w / y+Y ; c(3)G^ccΔ2^/TM3, Sb*; *sv^spa-pol^*stock was generously provided by Dr. Katherine Billmyre. The presence of mutant alleles was verified where possible using allele-specific PCRs optimized for this purpose. Primer sequences are shown in Supplementary Table S2.

### Fly crosses

Flies that were *Oregon-R* and *net dpp^d-ho^ dp b pr cn* were isogenized, then incorporated into various mutant backgrounds. The following stocks were built for this study: *y mei-218^1^/FM7* ; *net-cn* iso/*CyO, mei-218^6^ f / FM7* ; *OR+* iso*/CyO, net-cn* iso*/CyO* ; *rec^1^ Sb/TM6B Hu Tb, OR+* iso*/CyO* ; *kar ry^606^ rec^2^/MKRS Sb, OR+* iso*/CyO* ; *Su(var)3-9^06^/MKRS, Sb, net-cn* iso*/CyO* ; *Su(var)3-9^17^/MKRS Sb, y w* ; *OR+* iso*/CyO* ; *c(3)G^ccΔ2^/MKRS, y w* ; *net-cn iso/CyO* ; *c(3)G^ccΔ2^/TM6B*.

### Recombination mapping

Meiotic crossovers were mapped on chromosome *2* by crossing females that were heterozygous for the markers *net dpp^d-ho^ dp b pr* and *cn* in the mutant background of choice to males homozygous for the same markers. Mitotic crossovers were mapped by crossing males that were heterozygous for these markers on chromosome *2* and were *Su(var)3-9^06^/+* or *Su(var)3-9^06^/Su(var)3-9^17^* chromosome *3* to females homozygous for the chromosome *2* markers. Males and females were both between 1 and 5 days old when mated, and each vial was flipped after seven days. Progeny were scored for all phenotypic markers and any that had a pericentromeric crossover (between *pr* and *cn*) were collected to fine-map where within the pericentromere the crossover occurred, through allele-specific PCR. Complete datasets for all recombination mapping are given in Supplementary Table S1. Wild-type crossover distributions were taken from a previous recombination mapping dataset (PAZHAYAM *et al*. 2023). Total chromosome *2* crossover numbers for wild type were estimated using the same dataset, based on total proximal crossovers collected in this study (n=132), and is indicated as “adjusted total crossovers” in Supplementary Table S1. For *c(3)G^ccΔ2^,* fine-mapping of pericentromeric crossovers was done in 171 of the 478 flies with pericentromeric crossovers, requiring an adjusted total crossover number for percentages of total crossovers calculated in Table 1. This adjusted total crossover number is also indicated in Supplementary Table S1.

### Recombination calculations

Genetic length was calculated in centiMorgans (cM) as follows: (r/n) * 100, where *r* represents the number of recombinant flies in an interval (including single, double, and triple crossovers) and *n* represents total flies that were scored for that genotype. Release 6.53 of the reference genome of *Drosophila* was used to calculate physical length between chromosome *2* markers used for phenotypic recombination mapping. Since alpha heterochromatin sequence is not yet assembled, we estimated the length from the estimated heterochromatic sequence, 5.4 Mb for *2L* and 11.0 Mb for *2R* (Adams *et al*. 2000), minus the length of beta heterochromatin sequence in the Release 6.53 assembly (1.39 Mb for *2L*, 7.6 Mb for *2R*). CE values were calculated as 1-(observed crossovers/expected crossovers). Expected crossovers = total crossovers in a genotype * (physical length of proximal interval/total physical length).

### SNPs defining pericentromeric regions

Illumina sequencing was done on isogenized stocks of *Oregon-R* and *net-cn* to identify SNP differences. DNA from ∼50 whole flies was extracted using the QIAGEN DNeasy Blood and Tissue Kit and sequenced on the Illumina NovaSeq 6000. Reads were aligned to the reference genome using bowtie2 (v2.5.3) (Langmead and Salzberg 2012) and PCR and optical duplicates were marked using samtools markdup (v1.21) (Danecek *et al*. 2021). Variants were called using freebayes (v1.1.0) (Erik Garrisson 2012). Unique SNPs between the *net-cn* and *OR*+ chromosome *2* were identified using bcftools isec (v1.20) (Danecek *et al*. 2021). SNPs were validated by analyzing reads using Integrative Genomics Viewer (Robinson *et al*. 2011) and via PCR.

Four SNPs (called *beta2L*, *alpha2L*, *alpha2R*, and *beta2R*) were chosen to mark the boundaries between proximal euchromatin, beta heterochromatin, and alpha heterochromatin on each arm of chromosome *2*. The *alpha2L* (position 23424573, C in *net-cn*, A in *OR+*) and *alpha2R* (position 639629, C in *net-cn*, A in *OR+*) SNPs chosen were the most proximal chromosome *2* SNPs in (Hartmann *et al*. 2019b). The *beta2L* (position 22036096, A in *net-cn*, T in *OR+*) and *beta2R* (position 5725487, C in *net-cn*, T in *OR+*) SNPs chosen were based on maximum proximity to the heterochromatin-euchromatin boundary as defined by various studies summarized in Supplemental Table S3 of (Stutzman *et al*. 2024).

Proximal euchromatin is defined as the region between phenotypic marker *pr* and the *beta2L* SNP on chromosome *2L* and the region between the *beta2R* SNP and the phenotypic marker *cn* on chromosome *2R*. Beta heterochromatin is defined as the region between the *beta2L* SNP and *alpha2L* SNP on chromosome *2L* and the *alpha2R* SNP and *beta2R* SNP on chromosome *2R.* Alpha heterochromatin is defined as the region between the *alpha2L* SNP on chromosome *2L* and the *alpha2R* SNP on chromosome *2R*.

A second *beta2R* SNP (position 5726083, A in *net-cn*, T in *OR+*) was chosen for the progeny of *Su(var)3-9^06^*/+ and *c(3)G^ccΔ2^* mutants with pericentromeric crossovers as the allele-specific PCR amplifying the *beta2R* SNP at position 5725487 was no longer robust towards the end of our study. For consistency, progeny of WT, *mei-218^null^*, *rec^null^*, and *Su(var)3-9^06^/Su(var)3-9^17^* flies with pericentromeric crossovers where the position of the crossover was indicated by the presence or absence of the 5725487 *beta2R* band were re-confirmed with the allele-specific PCR amplifying the *beta2R* SNP at position 5726083. Additional SNPs *alpha2L_II* (position 23423662, A in *net-cn*, C in *OR+*) and *alpha2R_II* (position 637775, T in *net-cn*, C in *OR+*) were used to confirm each alpha-heterochromatic crossover that was observed. Primer sequences and PCR conditions are shown in Supplementary Table S3. Optimization PCRs for each SNP are shown in Supplementary Figure S2.

### Allele-specific PCR

Progeny from the crosses of experimental females of the desired mutant background and males homozygous for phenotypic markers *net-cn* that had a pericentromeric crossover (a crossover between the most proximal markers *purple* and *cinnabar* on either arm of chromosome *2*) were collected and DNA was extracted. Since the recombined chromosome from experimental females is recovered over a *net-cn* chromosome from males, all progeny carry the *net-cn* versions of each SNP. Therefore, allele-specific PCRs that amplify the *OR+* versions had to be performed on progeny with a pericentromeric crossover to map whether the crossover occurred in proximal euchromatin, beta heterochromatin, or alpha heterochromatin. For each allele-specific PCR, the presence of a band indicates that the recombined chromosome from the experimental female has the *OR+* version of the SNP. The absence of a band indicates that the recombined chromosome from the experimental female has the *net-cn* version of the SNP. With this information, we pinpointed the switch from *OR+* SNPs to *net-cn* SNPs on the recombined chromosome, telling us where the pericentromeric crossover in the experimental female occurred. Gels from all allele-specific PCRs for each fly of every genotype (WT, *mei-218^null^*, *rec^null^*, and *Su(var)3-9^06^/Su(var)3-9^17^, Su(var)3-9^06^*/+ and *c(3)G^ccΔ2^*) are shown in Supplementary Figure S1.

## Data Availability Statement

*Drosophila* stocks are available upon request. The authors confirm that all data necessary for confirming the conclusions of the article are present within the article, figures, table, and supplemental information. Illumina sequences for isogenized *OR*+ and *net-cn* flies have been submitted to SRA under BioProject PRJNA1198609.

## Acknowledgments

We thank Katie Billmyre and Gary Karpen for generously sharing *Drosophila* stocks, Carolyn Turcotte for help processing Illumina sequences/SNP calling, and Greg Copenhaver, Kacy Gordon, Dale Ramsden, Dan McKay, as well as members of the Sekelsky lab for helpful comments.

## Funding

This work was supported by a grant from the National Institute of General Medical Sciences to JS under award 1R35GM118127. NP was supported in part by a grant from the National Institute on Aging under award 1F31AG079626 and the National Institute of General Medical Sciences under award T32GM135128.

## Competing Interests

The authors declare that they have no conflicts of interest.

---

**<u>Figure S1.</u>** Gel images for allele-specific PCRs used to amplify the *OR+* version of each SNP that defines the various pericentromeric regions in WT as well as each mutant in this study. Numbers indicate each fly with a pericentromeric crossover that was analyzed. O indicates *OR*+ flies (positive control) and N indicates *net-cn* flies (negative control).

**<u>Figure S2.</u>** Gel images for optimization PCRs performed to decide on the conditions for allele-specific PCRs that were used to amplify the *OR+* version of each SNP that defines the various pericentromeric regions in this study. Temperature gradients from 56C to 66C are indicated on each gel. A black box is drawn around the band that should be amplified in *OR*+ (O) flies and not in *net-cn* (N) flies.

**<u>Table S1.</u>** The complete meiotic crossover distribution dataset on chromosome *2* between markers *net* and *cinnabar* for wild type and each mutant in this study. Mitotic crossover distribution datasets between the same chromosome *2* markers for *Su(var)3-9^06^/+* and *Su(var)3-9^06^/Su(var)3-9^17^* flies are also shown. SCO, DCO, and TCO denote single, double, and triple crossover progeny, respectively.

**<u>Table S2.</u>** Primer sequences used to validate various mutant alleles.

**<u>Table S3.</u>** Primer sequences and PCR conditions for allele-specific PCRs used to amplify the *OR+* version of each SNP that defines the various pericentromeric regions in this study.

